# FRIZZLED 2 regulates limb development by mediating both β-catenin-dependent and independent Wnt signaling pathways

**DOI:** 10.1101/2022.09.08.507207

**Authors:** Xuming Zhu, Mingang Xu, N. Adrian Leu, Edward E. Morrisey, Sarah E. Millar

## Abstract

Human Robinow Syndrome and omodysplasia, characterized by skeletal limb and craniofacial defects, are associated with mutations in the Wnt receptor FZD2. However, as FZD2 can activate both canonical and non-canonical Wnt pathways, its precise functions and mechanisms of action in limb development are unclear. To address these questions, we generated mice harboring a single nucleotide insertion in the Dishevelled-interacting domain of *Fzd2* (*Fzd2^em1Smill^*), causing a frameshift mutation similar to the effects of human syndromic *FZD2* mutations. *Fzd2^em1Smill^* mutant mice had shortened limbs resembling those of Robinow Syndrome and omodysplasia patients. *Fzd2^em1Smill^* mutant embryos displayed decreased canonical Wnt signaling in developing limb mesenchyme and disruption of digit chondrocyte elongation and orientation, which is controlled by the WNT5A/PCP pathway. In line with this, we found that tissue-specific disruption of *Fzd2* function in limb mesenchyme caused formation of shortened bone elements and was associated with deficiency in both Wnt/β-catenin and WNT5A/PCP signaling. These findings indicate that FZD2 controls limb development by mediating both canonical and non-canonical Wnt pathways and reveal causality of pathogenic *FZD2* mutations in Robinow Syndrome and omodysplasia patients.

**Summary statement:** Human FZD2 mutations are associated with limb defects; using genetic mouse models we revealed causality of these mutations and showed that they disrupt both canonical and non-canonical Wnt signaling.

## Introduction

Congenital limb defects (CLDs) are common in human populations (Ephraim et al., 2003). Genetic defects are a major cause of CLDs, and have been characterized by next generation sequencing (Carli et al., 2013; White et al., 2018). While the genetic defects causing CLDs are quite diverse, many of these affect specific signaling pathways. For example, mutations in Wnt/PCP pathway components have been identified in patients suffering from Robinow syndrome, which is characterized by shortened limbs as well as craniofacial defects (White et al., 2018).

Wnt signaling pathways play key roles in many developmental processes and human diseases (Klaus and Birchmeier, 2008). Activation of the canonical pathway is initiated by the binding of WNT ligands such as WNT1 and WNT3 to a specific subset of FRIZZLED (FZD) receptors, causing stabilization of cytoplasmic β-catenin and its translocation to the nucleus where it activates transcription of downstream target genes such as *Axin2* (MacDonald et al., 2009). Non-canonical Wnt pathways are triggered by WNT5A and WNT11, which regulate cytoskeletal arrangement, cell orientation, and cell movements via the planar cell polarity (PCP) or Ca^2+^ pathways (Gordon and Nusse, 2006).

In vertebrates, at least fifteen Wnt family members are expressed in developing limbs (Witte et al., 2009). Multiple lines of evidence demonstrate that Wnt/β-catenin signaling is essential for limb development. For instance, deletion of *Wnt3* or *Ctnnb1* in mouse limb ectoderm causes failure of normal formation of the apical ectodermal ridge (AER) and limb agenesis (Barrow et al., 2003). Consistent with this, mutations of human *WNT3* are associated with tetra-amelia syndrome, characterized by severe defects in limb development (Niemann et al., 2004). Specific deletion of β-catenin in limb mesenchyme disrupts AER integrity and induces apoptosis of mesenchymal cells (Hill et al., 2006). Mouse genetic studies have also revealed critical roles for WNT/PCP signaling in limb development: for instance, deletion of the *Wnt5a, Ror2* or *Vangl2* genes prevents proper elongation and orientation of differentiating chondrocytes along the proximal-distal (PD) axis of the limb (Gao et al., 2011; Yamaguchi et al., 1999). *WNT5A* and *ROR2* mutations are associated with Robinow Syndrome in human patients, who display a phenotype of shortened, but normally patterned, limbs similar to that observed in *Wnt5a* and *Ror2* loss of function mutant mice (White et al., 2018).

Although the roles of WNT ligands and their downstream pathways in limb development have been intensively investigated, the functions of specific FZD transmembrane receptors in this process are less clear. FZD proteins directly bind WNT ligands, and their intracellular C-terminal domains interact with Dishevelled (DVL) to transduce canonical or non-canonical Wnt signaling (Gordon and Nusse, 2006). Accumulating evidence suggests that activation of canonical versus non-canonical signaling by FZD receptors depends on their binding to specific WNT ligands. For example, FZD2 interacts with WNT3 and WNT3A to stabilize β-catenin and activate the canonical Wnt pathway in a myeloid progenitor cell line (Dijksterhuis et al., 2015). By contrast, FZD2 interacts with WNT5A to mediate non-canonical Wnt signaling and stimulate epithelial-mesenchymal transition (EMT) and migration in a variety of tumor cell lines (Gujral et al., 2014). Human genetic studies have identified heterozygous *FZD2* mutations in Robinow Syndrome and omodysplasia patients. These include missense or dinucleotide substitution mutations that remove Gly434, a residue that is conserved in all vertebrates and is located at the edge of a transmembrane domain next to the DVL interacting domains, and nonsense mutations that result in truncation of all the DVL interacting domains, or affect the function of the last DVL interacting domain (Nagasaki et al., 2018; Saal et al., 2015; Turkmen et al., 2017; Warren et al., 2018; White et al., 2018). These data suggest involvement of FZD2 in regulating limb development and specifically implicate importance of the DVL interacting domains. However, this hypothesis has not been tested through loss of function studies in a genetically manipulable model system. Furthermore, whether *Fzd2* activates canonical or non-canonical signaling during limb development is unclear.

To address these questions, we generated mice harboring a single nucleotide insertion in the Dishevelled-interacting domain of *Fzd2 (Fzd2^em1Smill^*, hereafter referred to as *Fzd2^INS^*), causing a frameshift mutation similar to the effects of human syndromic *FZD2* mutations. *Fzd2^INS^* mice mimicked the phenotypes seen in Robinow Syndrome and omodysplasia patients indicating that *FZD2* mutations underlie these conditions. We also utilized conditional loss of function *Fzd2^fl^* mice together with *Prx1-Cre* that is active in limb mesenchyme. We found that *Fzd2* deficiency in limb mesenchyme caused formation of shortened bone elements and was associated with disruption of both Wnt/β-catenin and WNT5A/PCP signaling. These findings demonstrate that FZD2 controls limb development by mediating different Wnt signaling pathways.

## Results

### *Fzd2* is ubiquitously expressed in the ectoderm and mesenchyme of developing limb buds

To determine the location of *Fzd2* expression in developing mouse limb buds, we carried out in situ hybridization (ISH) for *Fzd2* mRNA at embryonic day (E) 9.5 and E10.5. At E9.5, *Fzd2* is ubiquitously expressed in both ectoderm and mesenchyme of the emerging forelimb bud, with lower levels of expression in the ectoderm and in mesenchyme immediately underlying the AER, and most intense expression in more proximal mesenchymal cells (Fig. 1A-A’). This expression pattern persists in the forelimb bud at E10.5 (Fig.1B-B’), and is similar in the E10.5 hindlimb bud (Fig. 1C-C’).

**Figure 1.**
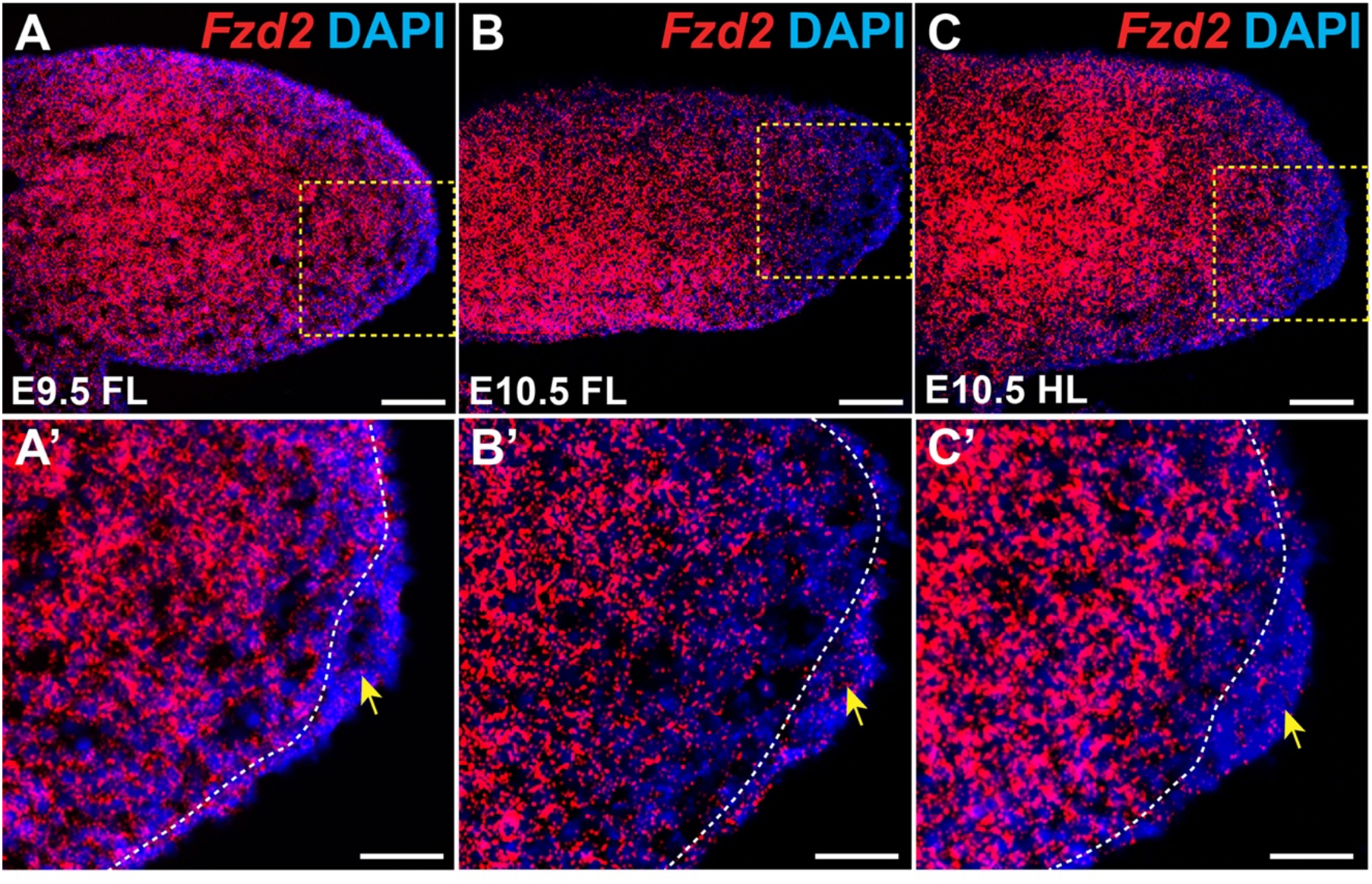
*Fzd2* is broadly expressed in the developing limb bud. (A) RNAscope ISH of sagittal sections shows that *Fzd2* is expressed in forelimb bud at E9.5. (A’) Magnified view of the indicated region in panel (A). *Fzd2* expression is higher in the mesenchyme than in the ectoderm, including the AER. (B) *Fzd2* expression persists in the ectoderm and mesenchyme of the forelimb bud at E10.5 but is decreased in distal mesenchyme compared with its expression at E9.5. (B’) Magnified view of the indicated region in (B). (C) *Fzd2* expression in E10.5 hindlimb is similar to that in the forelimb. (C’) Magnified view of the indicated region in (C). In all photomicrographs, dorsal limb is oriented at the top and distal to the right. Yellow arrows indicate AER; white dashed lines mark the boundary between ectoderm and mesenchyme. N=3 samples analyzed per stage. FL, forelimb; HL, hindlimb. Scale bars: panels (A-C), 100μm; panels (A’-C’), 50μm.

### An insertional mutation in the DVL-interacting domain of FZD2 causes lethality of pups after birth

To determine the functional consequences of the C-terminal FZD2 mutations observed in Robinow Syndrome and omodysplasia patients, we used CRISPR/Cas9 gene editing to generate a disruption in mouse *Fzd2* similar to those observed in human patients. The modified *Fzd2* allele (*Fzd2^INS^*) harbors an extra guanine between c.1656 and c.1657 (NM_020510.2: c.1656_1657insG), leading to a frame-shift mutation that mutates a DVL binding motif (KTxxxW) and removes the PDZ-interacting domain (ETTV), resulting in an aberrant C-terminus and a predicted protein that is 39 amino acids longer than wild-type FZD2 (Fig. 2A). *Fzd2^INS/+^* mice lacked overt phenotypes and were fertile. However, *Fzd2^INS/INS^* pups died within a few days of delivery, and had extended abdomens containing air but little milk, suggesting difficulty in feeding (Fig. 2B). *Fzd2* mRNA and protein levels were comparable between control and *Fzd2^INS/INS^* samples (Fig. 2C). Detailed analysis revealed that the *Fzd2^INS/INS^* pups had a 100% penetrant phenotype of cleft palate and reduced distances from the back of the skull to the anterior tip of the upper jaw (n=3 mutants and n=4 controls analyzed; p=0.001) (Fig. 2D). These results were consistent with a prior study demonstrating a 50% penetrant cleft palate phenotype in mice carrying a hypomorphic mutation in *Fzd2* (Michalski et al., 2021; Yu et al., 2010).

**Figure 2.**
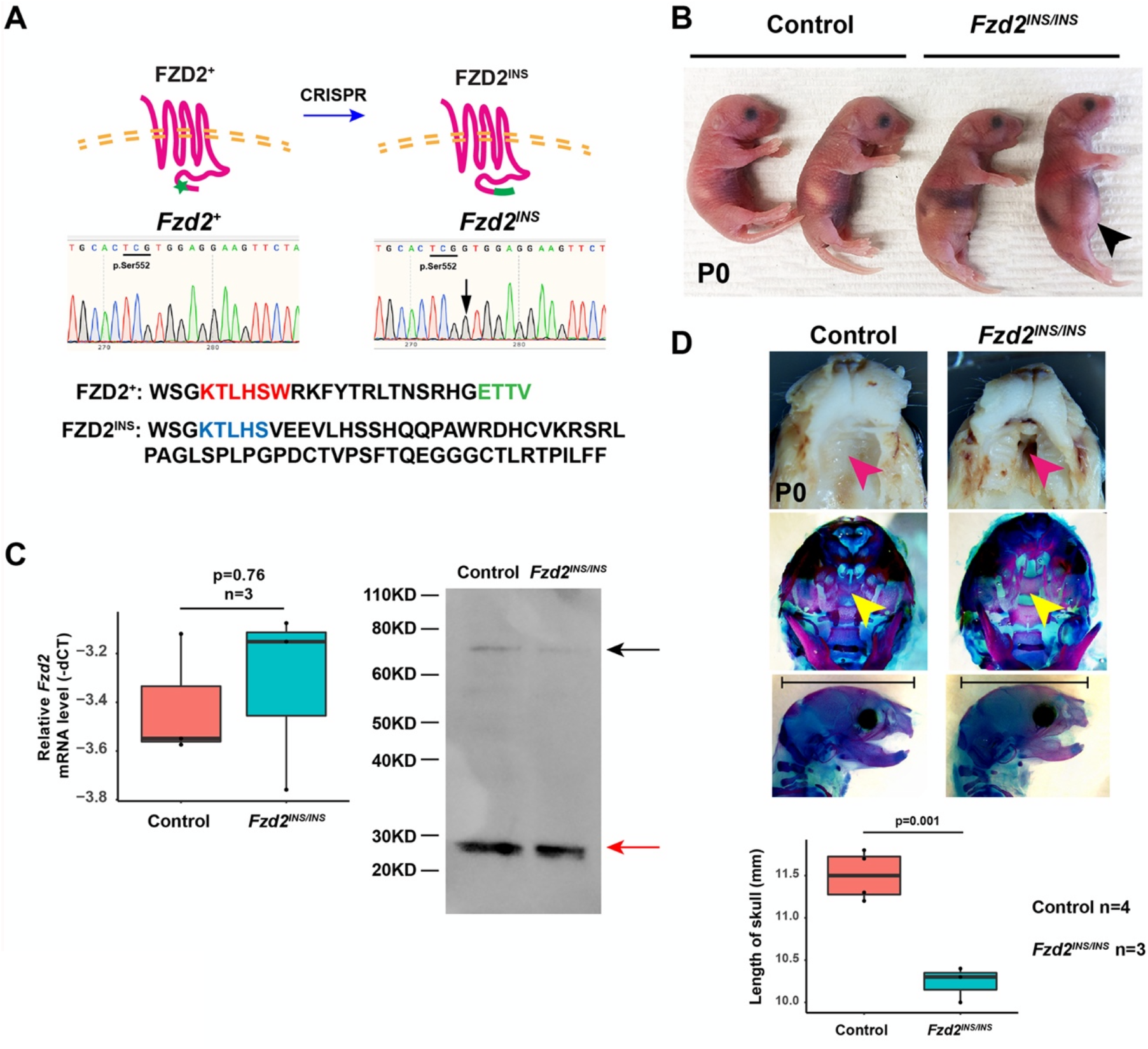
*Fzd2^INS/INS^* pups display craniofacial defects. (A) CRISPR/Cas9 editing introduced an extra guanine (G) directly after p.Ser552, mutating the DVL binding motif (red type), removing the PDZ-interacting domain (green type), and causing a frameshift in the C-terminus of FZD2. (B) *Fzd2^INS/INS^* neonates had swollen abdomens lacking milk spots (black arrowhead). (C) qPCR and immunoblotting showed that *Fzd2* mRNA and protein levels were similar in control and *Fzd2^INS/INS^* skin samples. The black arrow indicates FZD2 protein; the red arrow indicates a nonspecific band that serves as a loading control. (D) *Fzd2^INS/INS^* pups display severely clefted palates (pink and yellow arrowheads) and shortened skulls. The lengths of four control and three mutant skulls were measured. Two-tail Student’s *t*-test was used to calculate p-value. p<0.05 was considered to be significant.

### *Fzd2^INS/INS^* mice show abnormal limb development

*Fzd2^INS/INS^* pups did not show obvious limb patterning defects but displayed statistically significantly shorter limb bones than littermate controls (Fig. 3A and 3B). To determine the mechanisms underlying limb defects in *Fzd2^INS/INS^* mice, we analyzed the effects of the *Fzd2^INS^* mutation on canonical and non-canonical Wnt signaling pathways. mRNA levels for *Wnt3*, which is expressed in limb ectoderm and directs canonical β-catenin-mediated signaling, were unaffected by the *Fzd2^INS^* mutation at E12.5; however, levels of canonical signaling, indicated by expression of the ubiquitous canonical Wnt target gene *Axin2*, were downregulated in the mesenchyme of *Fzd2^INS/INS^* mutants compared with littermate controls (Fig. 3C), indicating that FZD2 mutation disrupts canonical signaling downstream of WNT3.

**Figure 3.**
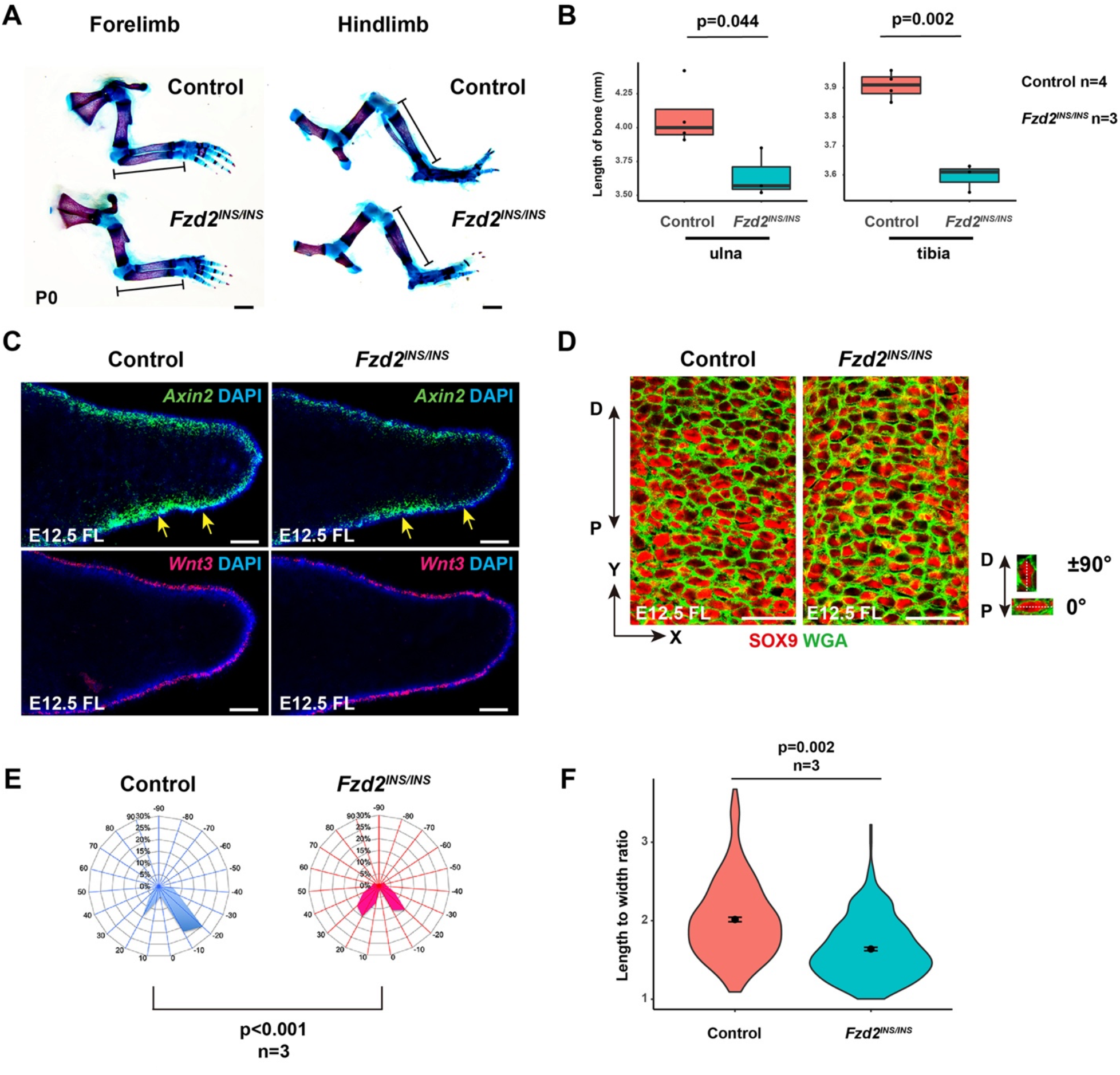
The *FZD2^INS^* mutation causes defective limb development by altering both canonical Wnt and Wnt/PCP signaling. (A) Skeletal preparations show that the ulna and tibia of *Fzd2^INS/INS^* pups are shorter than those of littermate controls at P0. (B) Quantitation of bone element lengths in *Fzd2^INS/INS^* pups (n=4 control and 3 mutants). P-values were calculated with two-tailed Student’s *t*-test. p<0.05 was considered to be significant. (C) *Fzd2* deficiency causes reduced expression of *Axin2* (yellow arrows) but has little effect on *Wnt3* expression in forelimbs at E12.5. (D) WGA and SOX9 staining show that elongation and orientation of digit chondrocytes is affected in *Fzd2^INS/INS^* mutants. (E) Quantitation of cell orientation in E12.5 forelimbs. One hundred chondrocytes from each embryo were measured. Three pairs of control and mutant embryos were analyzed. The X-axis represents the angle of orientation; the Y-axis represents the percentage of chondrocytes at angle X. Chondrocytes oriented horizontally are designated as 0°; and chondrocytes oriented along the P-D axis are designated as ± 90°. The Kolmogorov Smirnov test was used to calculate the p-value. p<0.05 was considered to be significant. (F) Quantitation of the ratio of length to width of chondrocytes shows that this is significantly altered in E12.5 *Fzd2^INS/INS^* mutant forelimbs. One hundred chondrocytes from each embryo and three embryos of each genotype were analyzed. Two-tailed Student’s *t*-test was used to calculate p-value. p<0.05 was considered to be significant. Scale bars: (A), 1mm; (C), 100μm; (D), 25μm.

To assay for effects on non-canonical Wnt signaling, we measured the elongation and orientation of SOX9-expressing chondrocytes in distal digits, which is controlled by the WNT5A/PCP pathway (Yang et al., 2017). In E12.5 control digits, the majority of distal chondrocytes were elongated, and their major axes trended to be perpendicular to the P-D axis of the limbs; however, in *Fzd2^INS/INS^* digits, the elongation and orientation of chondrocytes was abnormal (Fig. 3D-F), similar to limb chondrocyte phenotypes in *Wnt5a^-/-^* embryos (Gao et al., 2011). Taken together, these data indicate FZD2 mediates both canonical and WNT5A/PCP signaling pathways in limb development.

### Mutation of mesenchymal *Fzd2* produces severe limb defects

As *Fzd2* is mainly expressed in limb mesenchymal cells, we asked whether mesenchymal *Fzd2* is required for normal limb development by using *Prx1-Cre* to drive *Fzd2* mutation. We assessed the effects of loss of mesenchymal *Fzd2* function in developing forelimb buds because *Prx1-Cre* drives mosaic recombination in hindlimb buds, which complicates analysis (Logan et al., 2002). We utilized *Fzd2^fl/fl^* mice that permit conditional *Fzd2* mutation (Kadzik et al., 2014). Although originally described as a conventional floxed allele, a subsequent study (Michalski et al., 2021) and our data show that this *Fzd2^fl^* allele contains an inverted duplication of *Fzd2* with oppositely oriented *loxP* sites positioned respectively in the 3’ UTR and 5’ UTR of the duplicated genes (Fig. S1A). Upon Cre mediated recombination, the sequence between the two *loxP* sites undergoes continuous inversion (Michalski et al., 2021) (Fig. S1B). This results in production of an inverted transcript complementary to *Fzd2* mRNA that is likely to function as an antisense RNA (Fig. S1B and C) and could explain loss of FZD2 function in *Fzd2^fl/fl^* mice that express Cre recombinase.

We found that *Prx1-Cre Fzd2^fl/+^* mutants were viable and fertile. Their forelimbs had normal digits but were shortened. More severe forelimb hypoplasia, with loss of almost all bone elements, was observed in postnatal *Prx1-Cre Fzd2^fl/fl^* mutants (Fig. 4A, B). Similar phenotypes were observed at E17.5 (Fig. 4C-H). ISH experiments confirmed that mesenchymal *Fzd2* mRNA levels were reduced in *Prx1-Cre Fzd2^fl/+^* and almost absent in *Prx1-Cre Fzd2^fl/fl^* forelimb mesenchyme during embryogenesis (Fig. 5A-C). These data demonstrate that mesenchymal *Fzd2* regulates limb development in a dose-dependent manner.

**Figure 4.**
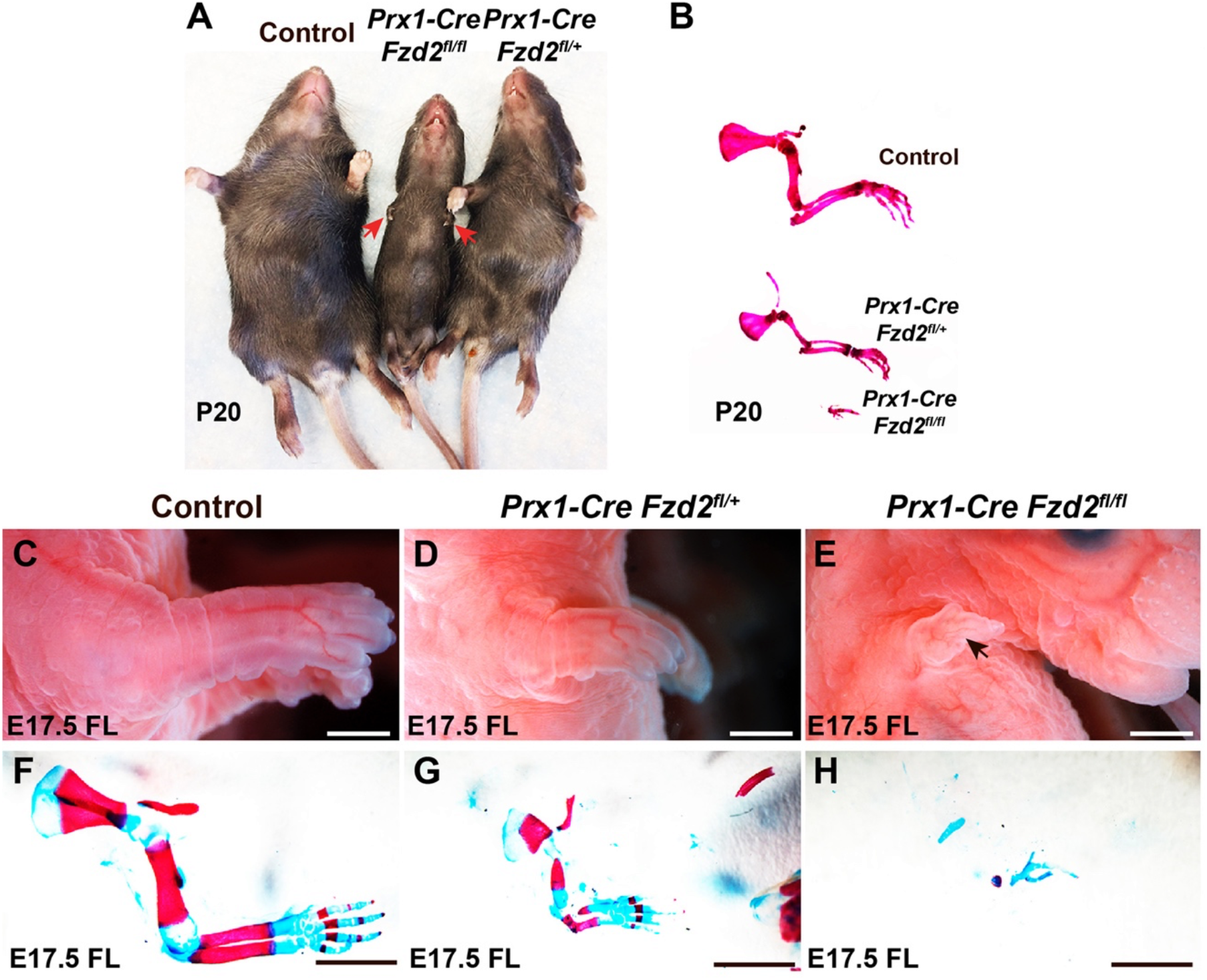
Mesenchymal *Fzd2* is required for normal limb development. (A) *Prx1-Cre Fzd2^fl/+^* and *Prx1-Cre Fzd2^fl/fl^* mutants have hypoplastic forelimbs at P20. (B) Skeletal preparation of P20 forelimbs. Compared with control forelimb, *Prx1-Cre Fzd2^fl/+^* pups have hypoplastic forelimbs; *Prx1-Cre Fzd2^fl/fl^* pups only have a few residual bone elements in their forelimb. (C-E) Whole mount views of E17.5 control (C), *Prx1-Cre Fzd2^fl/+^* (D) and *Prx1-Cre Fzd2^fl/fl^* (E) forelimbs show reduced forelimb size in the heterozygous mutant (D) and a small residual forelimb in the homozygous mutant (E). (F-H) Skeletal preparations show that all the skeletal elements are hypomorphic in an E17.5 *Prx1-Cre Fzd2^fl/+^* heterozygous mutant forelimb (G) and only a few skeletal elements develop in a *Prx1-Cre Fzd2^fl/fl^* homozygous mutant (H) compared with control (F). Scale bars: 0.5mm.

**Figure 5.**
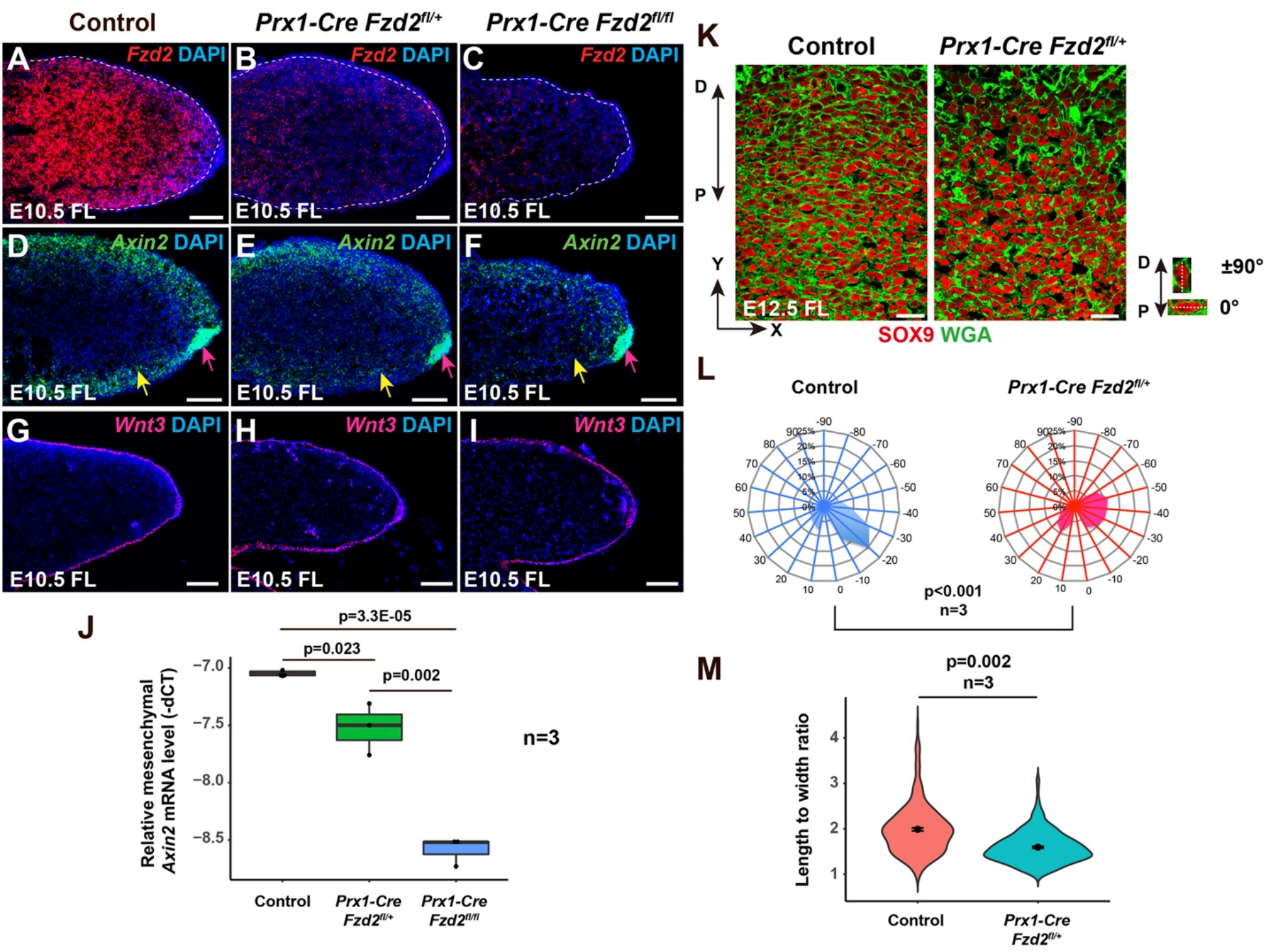
Reduced mesenchymal *Fzd2* expression affects both canonical and non-canonical Wnt signaling. (A-C) Mesenchymal *Fzd2* expression in E10.5 forelimb mesenchyme is reduced in *Prx1-Cre Fzd2^fl/+^* heterozygous mutants (B) and almost absent in *Prx1-Cre Fzd2^fl/fl^* homozygous mutants (C) compared with control (A). (D-F) ISH shows that *Axin2* expression is reduced in distal forelimb mesenchyme (yellow arrows) but not in the AER (pink arrows) in E10.5 *Prx1-Cre Fzd2^fl/+^* (E) and *Prx1-Cre Fzd2^fl/fl^* (F) embryos compared with control (D). (G-I) Ectodermal *Wnt3* expression is unaffected in the forelimb of E10.5 *Prx1-Cre Fzd2^fl/+^* (H) and *Prx1-Cre Fzd2^fl/fl^* (I) embryos compared with control (G). (J) qPCR quantification of *Axin2* mRNA levels in E10.5 forelimb mesenchyme shows statistically significantly decreased *Axin2* expression upon mesenchymal *Fzd2* mutation. n=3 per genotype. Two-tailed Student’s *t*-test was used to calculate p-values. p<0.05 was considered to be significant. (K) WGA (green) stains cell membranes revealing the shapes of SOX9-expressing chondrocytes. In E12.5 control forelimb digits, chondrocytes are elongated and predominantly lie perpendicular to the P-D axis; in *Prx1-Cre Fzd2^fl/+^* digits, the chondrocytes are more rounded and their orientation is more random. (L) Quantitation of chondrocyte orientation. At least 500 control and *Prx1-Cre Fzd2^fl/+^* chondrocytes from three embryos of each genotype were analyzed. The X-axis represents the angle of orientation; the Y-axis represents the percentage of chondrocytes at each angle X. Chondrocytes oriented horizontally are designated as 0°, and chondrocytes oriented along the P-D axis are designated as ± 90°. A Kolmogorov Smirnov test was used to calculate the p-value. p<0.05 was considered to be significant. (M) Quantitation of the length to width ratio of *Fzd2* deficient chondrocytes compared with controls shows a statistically significant difference, demonstrating that chondrocyte cell shape is altered in the mutants. One hundred chondrocytes from each embryo were analyzed from n=3 embryos per genotype. Two-tailed Student’s *t*-test was used to calculate the p-value. p<0.05 was considered to be significant. Scale bars: panels (A-I), 100μm; panel (K), 20μm.

### *Fzd2* mediates both canonical and non-canonical Wnt signaling in forelimb mesenchyme

As mesenchymal canonical Wnt/β-catenin signaling is required for limb development (Hill et al., 2006), we asked whether the Wnt/β-catenin pathway is affected in *Fzd2* deficient mesenchyme. Expression of *Axin2*, a ubiquitous target of canonical signaling, was attenuated in a dosedependent manner in limb bud mesenchyme, but not in the ectoderm, upon mesenchymal *Fzd2* mutation (Fig. 5D-F and J). Ectodermal *Wnt3* expression was not affected (Fig. 4G-I), indicating that mesenchymal FZD2 mediates β-catenin signaling in the mesenchyme downstream of ectodermal WNT3.

We tested whether the non-canonical Wnt pathway is also affected upon mesenchymal *Fzd2* mutation by assaying elongation and orientation of SOX9-expressing chondrocytes in the distal digits, which is controlled by WNT5A/PCP signaling (Yang et al., 2017). As most bone elements are missing in *Prx1-Cre Fzd2^fl/fl^* forelimbs, we assayed for digit chondrocyte elongation and orientation in E12.5 *Prx1-Cre Fzd2^fl/+^* forelimbs. These experiments showed that digit chondrocyte elongation and orientation was abnormal in *Prx1-Cre Fzd2^fl/+^* mutants compared with controls (Fig. 5K-M), similar to the phenotypes observed in *Wnt5a^-/-^* embryos (Gao et al., 2011) and in *Fzd2^INS/INS^* mutants. Taken together, these observations indicate that FZD2 mediates both Wnt/β-catenin and WNT5A/PCP pathways in embryonic forelimb mesenchyme.

## Discussion

Dominant *FZD2* mutations are associated with Robinow syndrome and omodysplasia, but whether these are causative, and the precise mechanisms by which FZD2 might act to control limb development, have been unclear. Here, we show that limb and craniofacial defects observed in these syndromes were phenocopied in mice carrying an insertional allele that, similar to human pathogenic mutations, disrupts an essential DVL interaction domain of FZD2. We also observed similar limb phenotypes in mice with a tissue-specific *Fzd2* loss of function mutation in limb mesenchyme. These data provide definitive evidence for causality of *FZD2* mutations.

In addition to *FZD2* mutations, mutations in the non-canonical Wnt pathway components *WNT5A* and *ROR2* are associated with Robinow syndrome, suggesting that disruption of a non-canonical WNT5A-FZD2-ROR2 pathway might underlie defective limb development (White et al., 2018). In line with this, we found that chondrocyte elongation and orientation, which are controlled by non-canonical Wnt signaling, were disrupted in limb bud mesenchyme of *Fzd2^INS^* mice and in mice with mesenchymal-specific *Fzd2* mutation. Thus, disruption of the non-canonical pathway can account at least in part for limb phenotypes in Robinow Syndrome.

Interestingly, in vitro experiments showed that FZD2 protein carrying a G538A mutation identified in omodysplasia patients was unable to transduce WNT3A-triggered canonical Wnt signaling (Saal et al., 2015), suggesting that impaired Wnt/β-catenin signaling might also contribute to defective limb development in these patients. Consistent with this, we observed decreased canonical Wnt signaling in the limb buds of *Fzd2^INS/INS^* embryos.

In line with these observations, we found that *Prx1-Cre Fzd2^fl/+^* mice displayed limb shortening similar to that seen in Robinow syndrome and omodysplasia patients. Limb bone shortening was much more severe in homozygous *Prx1-Cre Fzd2^fl/fl^* mice, and both mutants exhibited decreased canonical Wnt signaling activity in limb bud mesenchyme. These effects were also observed when β-catenin is deleted using the same Cre line (Hill et al., 2006), suggesting that mesenchymal FZD2 regulates limb development in part via canonical Wnt signaling. In addition, *Prx1-Cre Fzd2^fl/+^* mice showed striking defects in chondrocyte elongation and polarization, indicating that non-canonical Wnt signaling was also disrupted in limb bud mesenchyme upon loss of mesenchymal *Fzd2* function.

Taken together, these data suggest that shortened limbs observed in Robinow syndrome and omodysplasia patients result from FZD2 deficiency in developing limb bud mesenchyme and are caused by defects in both canonical and non-canonical Wnt signaling.

We noted that, by contrast with Robinow syndrome patients and mice with mesenchymal *Fzd2* mutation, mice heterozygous for *Fzd2^INS^* did not display obvious limb defects and aberrant canonical and non-canonical Wnt signaling: these were only apparent in homozygous *Fzd2^INS/INS^* mutants. These observations suggest that the *Fzd2^INS^* is hypomorphic, as might be predicted since it only affects one of the three DVL-interacting domains in FZD2. In line with this, mice homozygous for a global *Fzd2* knockout allele produced by the International Mouse Phenotyping Consortium exhibit early embryonic lethality (Dickinson et al., 2016) (Michalski et al., 2021).

In summary, while at least seven *Fzd* genes are expressed in the developing mouse limb bud (Summerhurst et al., 2008), the functions of individual FZD family members in this context have been unclear. Our data identify FZD2 as a key component of both canonical and non-canonical Wnt signaling pathways in limb development and provide a mechanistic understanding of the defects in this process that are observed in patients carrying *FZD2* mutations. These findings will inform future research aimed at developing therapeutic interventions for Robinow Syndrome and omodysplasia patients.

## Materials and Methods

### Mice

The following mouse lines were used: *Fzd2^fl^* (Kadzik et al., 2014); *Prx1-Cre* (Jackson Laboratories, strain #005584); and *K14-Cre* (Jackson Laboratories, strain # 018964). All mice were maintained on a mixed strain background. Mice were allocated to experimental or control groups according to their genotypes, with control mice being included in each experiment. Male mice carrying *Prx1-Cre* were crossed with *Fzd2^fl/fl^* females to avoid potential germ line recombination. Investigators were not blinded during allocation and animal handling as information about genotype was required for appropriate allocation and handling. Immunostaining and ISH studies were carried out and data recorded in a blinded fashion. Up to five mice were maintained per cage in a specific pathogen free barrier facility on standard rodent laboratory chow (Purina, catalog #5001). All animal experiments were performed under approved animal protocols according to institutional guidelines established by the Icahn School of Medicine at Mount Sinai IACUC committee.

### Generation of CRISPR mutant mice

6–8-week-old female C57BL/6 mice were super-ovulated by IP injection of 5 IU of pregnant mare serum gonadotropin (PMSG) followed 48 hours later by 5 IU of human chorionic gonadotropin (hCG) and were mated to B6D2F1 males. A cocktail solution containing 50ng/μl gRNA (Integrated DNA Technologies) (5’-ACACTCGTCTCACCAACAGCCGG-3’) and 100ng/μl Cas9 mRNA (A29378, Thermo Fisher Scientific) was injected into one blastomere of two-cell embryos so that resulting embryos would be mosaic for the *Fzd2* mutation with at least 50% of the cells being wild-type. This approach was taken because injection into one-cell embryos was found to yield only homozygous mutants that were perinatal lethal and so could not be used to establish mutant lines. Embryos were incubated at 37°C, 5% CO_2_ in KSOM media (Millipore, MR-202P-5F). The KSOM culture drops were covered with mineral oil (Millipore, ES-005-C) to prevent evaporation. After cocktail injection, the 2-cell embryos were transferred the same day into the oviducts of Swiss Webster E0.5 pseudo-pregnant recipient females which were synchronized by using Swiss Webster vasectomized males. Micro-manipulation, embryo collection and embryo transfers were performed at room temperature in HEPES-buffered CZB medium. Founder mice were crossed with WT C57BL/6 mice to yield *Fzd2^INS/+^* offspring. *Fzd2^INS/+^* male and female mice were intercrossed to produce *Fzd2^INS/+^, Fzd2^INS/INS^*, and *Fzd2^+/+^* mice for analysis. Genotyping was performed by PCR using primers Fzd2^INS^-F: 5’-CACGACGGCACCAAGACGGA-3’, Fzd2^INS^-R: 5’-GAGACCGCTTCACACAGTG-3’. PCR product was then sequenced using primer Fzd2^INS^-F.

### RNA in situ hybridization

Embryos at the indicated stages were harvested and fixed with 4% paraformaldehyde (PFA) (Affymetrix/USB) in PBS overnight at 4°C. RNAscope was performed on fixed frozen sections following the user’s guide provided by Advanced Cell Diagnostic (ACD) and probes for *Fzd2* (ACD #565781), *Wnt3* (ACD #312241), and *Axin2* (ACD #400331). The sections were observed and photographed using a Leica Microsystems DM5500B fluorescent microscope.

### Skeletal preparation with Alcian blue and Alizarin red staining

Euthanized embryos or pups were skinned, eviscerated and fixed with 100% ethanol for 48 hours and transferred to acetone for 24 hours. The samples were placed in staining solution containing 0.015% Alcian blue and 0.005% Alizarin red for 1 week, and then treated with 1% KOH/10% glycerol until clear. Skeletal samples were photographed in 70% glycerol solution.

### Quantification of chondrocyte orientation and shape

E12.5 forelimbs were fixed in 4% PFA overnight at 4°C, embedded in OCT, and sectioned at 10μm. Sections were incubated with SOX9 antibody (Millipore, AB5535, 1:200) overnight at 4°C, followed by incubation with Alexa Fluor-labelled secondary antibodies (Thermo Fisher Scientific), and were washed with PBST. The sections were co-stained with Wheat Germ Agglutinin (WGA) according to the manufacturer’s instructions (Biotium). Sections were photographed using a Leica TCS SP8 confocal microscope (Leica Microsystems). The orientation of SOX9-expressing chondrocytes in the middle digit was determined by measuring the angle between the x-axis and the major axis of the chondrocyte. Chondrocyte shape was assayed by calculating the ratio of the lengths of the major axis and the minor axis. Data were analyzed by Image J (V1.49, NIH) and plotted using Microsoft Excel and R studio with ggplot2.

### Quantitative PCR

E10.5 limb buds were dissected and incubated in Dispase II (Gibco) solution for 30min at 37°C, and ectoderm was separated from the mesenchyme with forceps. Total RNA was extracted from limb bud mesenchyme using TRIzol (Thermo Fisher Scientific), purified using the RNeasy kit (Qiagen), and treated with the RNase-free DNase kit (Qiagen). Reverse-transcription was performed using a High-Capacity cDNA Reverse Transcription Kit (Applied Biosystems), and cDNA was subjected to real time PCR using the StepOnePlus system and SYBR Green Kit (Applied Biosystems). *Gapdh* was used as an internal control and expression differences were determined using the -ΔCT method. Primers for *Gapdh* were *Gapdh-F:* 5’-GAGAGGCCCTATCCCAACTC-3’, *Gapdh*-R: 5’-GTGGGTGCAGCGAACTTTAT-3’. Primers for *Axin2* were *Axin2*-F: 5’-GCTGGTTGTCACCTACTTTTTCTGT-3’, *Axin2*-R: 5’-GGGGAGCACTGTCTCGTCGTC-3’. Primers for *Fzd2* were *Fzd2*-F: 5’-CTTCACGGTCACCACCTATTT-3’, *Fzd2*-R: 5’-AACGAAGCCCGCAATGTA-3’.

### RT-PCR

Total RNA was extracted from keratinocytes of control and *K14-Cre Fzd2^fl/fl^* embryos at E14.5 using TRIzol (Thermo Fisher Scientific) and purified using the RNeasy kit (Qiagen). Total RNA samples were further treated with RNase-free DNase kit (Qiagen), reverse transcribed using a High-Capacity cDNA Reverse Transcription Kit (Applied Biosystems), and cDNA was subjected to PCR. The following primers were used: LoxP-F: 5’-GCCTGCTCGCTATTTTTGTTGGC-3’, LoxP-R: 5’-AAATGAGGAGGGAGAAAGAGGGGG-3’, *Fzd2*-R: 5’-AACGAAGCCCGCAATGTA-3’

### Western blotting

The back skin of neonatal pups was collected and lysed with RIPA buffer (Santa Cruz). Western blotting was performed using the XCell SureLock Mini-Cell Electrophoresis System (Invitrogen). FZD2 was detected by anti-FZD2 antibody (ab109094, Abcam, 1:500). Signal was detected and documented using a ChemiDoc MP system (Bio-Rad).

### Statistical analyses

Unpaired two-tailed Student’s t-test was used to calculate statistical significance for quantitation of bone element lengths and skull lengths; quantitation of ratios of chondrocyte length to width; and qPCR assay results. The Kolmogorov Smirnov test was used to calculate statistical significance for quantitation of chondrocyte orientation. p<0.05 was considered to be significant.

## Acknowledgements

We thank Dr. Kevin Kelley for CRISPR injections and Dr. Rachel Kadzik for helpful advice.

## Competing interests

The authors declare no competing interests.

## Funding

This work was supported by the National Institutes of Health 5R37AR047709 to S.E.M., the National Institutes of Health-funded University of Pennsylvania Skin Biology and Diseases Resource-based Center, P30AR069589 and the National Institutes of Health-funded Mount Sinai Skin Biology and Diseases Resource-based Center, P30AR079200.

## Data availability statement

The data that support the findings of this study are available from the corresponding author upon reasonable request.

## Author contributions

Xuming Zhu: conceptualization, formal analysis, investigation, visualization, writing – original draft preparation, writing – review & editing

Mingang Xu: conceptualization, formal analysis, investigation, visualization, writing – review & editing

Adrian Leu: investigation, methodology, writing – review & editing

Edward E. Morrisey: conceptualization, resources, writing – review & editing

Sarah E. Millar: conceptualization, funding acquisition, project administration, resources, supervision, visualization, writing – review & editing

## SUPPLEMENTARY INFORMATION

**Supplemental Figure S1.**
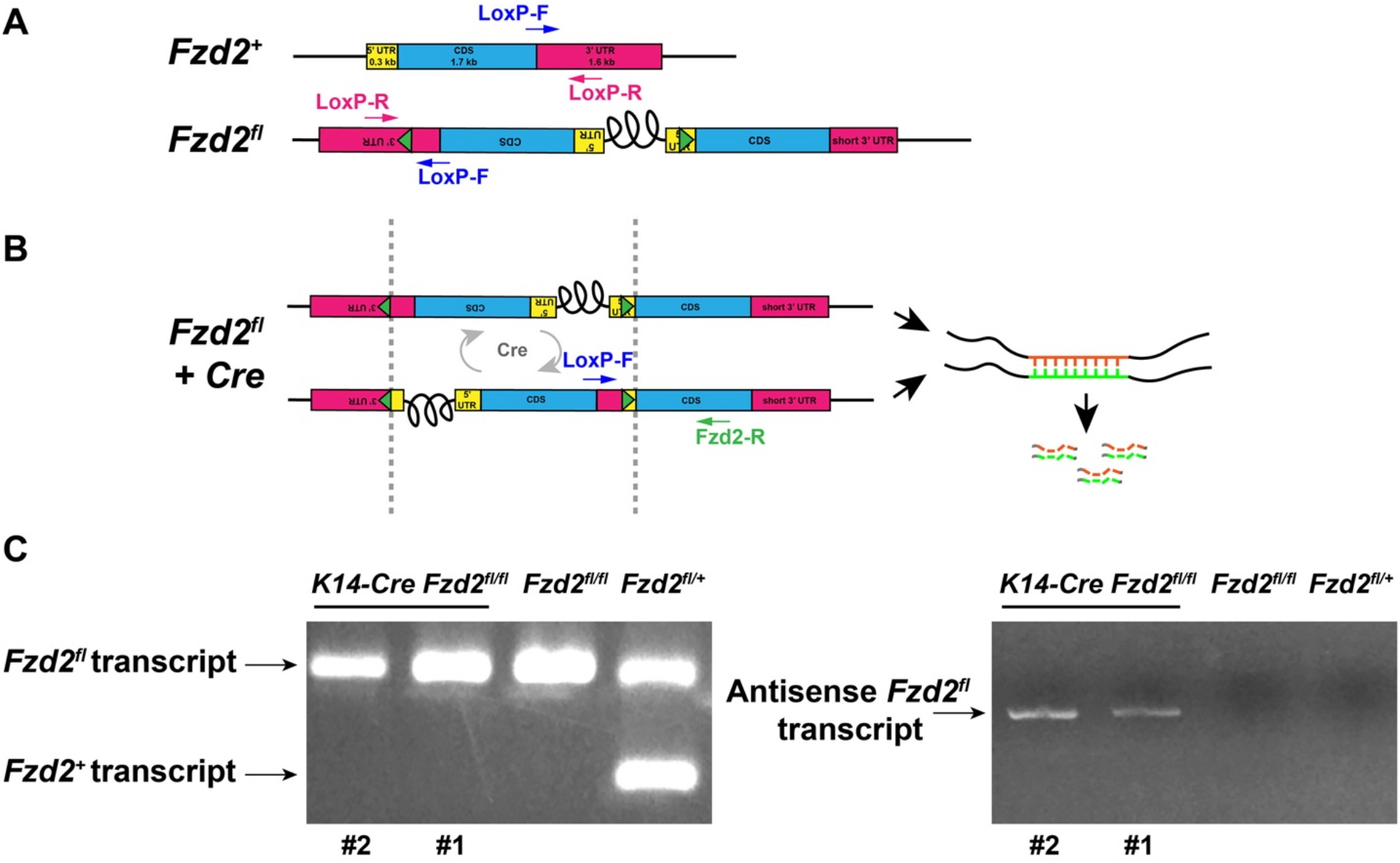
The *Fzd2^fl^* allele is a complex genomic alteration and generates antisense transcripts in the presence of Cre recombinase. (A) Schematic view of the wild type *Fzd2* allele (*Fzd2*^+^) and the *Fzd2^fl^* allele. (B) In the presence of Cre recombinase, the *Fzd2^fl^* allele is predicted to undergo continuous inversion, producing complementary mRNAs that could potentially bind to each other to prevent normal *Fzd2* translation. (C) In *Fzd2^fl/+^* keratinocytes, both *Fzd2^+^* and *Fzd2^fl^* mRNAs are detected by RT-PCR, whereas in *Fzd2^fl/fl^* keratinocytes, only *Fzd2^fl^* transcripts are detected (left panel). Antisense *Fzd2^fl^* transcripts are produced only in the presence of Cre recombinase (right panel).

## Notes

### Competing Interest Statement

The authors have declared no competing interest.

